# Downregulated expression of hepatic β-Klotho is associated with the hypertensive phenotype in SHR

**DOI:** 10.1101/2025.10.27.684772

**Authors:** Pedro Sousa Soares, Giovanna Arielle de Oliveira, Paula Magalhães Gomes, Vagner Roberto Antunes

## Abstract

**Background:** Fibroblast Growth Factor 21 (FGF-21) is an endocrine hormone that regulates metabolism and exerts cardiovascular effects through interaction with its co-receptor β-Klotho (KLB). Impaired FGF-21/KLB signaling has been linked to metabolic disorders, but its role in hypertension remains unclear.

**Objective:** This study investigated the expression of genes related to the FGF-21 signaling axis (FGF-21, FGFR1, and KLB) in the liver and central nervous system (CNS) of spontaneously hypertensive rats (SHR) at pre-hypertensive (21 days) and hypertensive (90 days) stages.

**Methods:** Systolic tail pressure (STP) and heart rate (HR) were recorded by tail-cuff plethysmography. Gene expression in the liver, brainstem, hypothalamus, and frontal cortex was quantified by RT-qPCR and normalized to β-actin.

**Results:** SHR-90d exhibited significantly higher STP compared to SHR-21d, with no difference in HR. Hepatic KLB mRNA expression was markedly reduced in SHR-90d compared to SHR-21d (p < 0.05), while FGF-21 and FGFR1 levels remained unchanged. No significant alterations in FGF-21 pathway genes were detected in CNS regions.

**Conclusion:** The selective downregulation of hepatic β-Klotho in hypertensive SHR suggests a state of peripheral FGF-21 resistance, potentially linking metabolic dysregulation to the maintenance of hypertension. These findings highlight the liver as a peripheral site coupling metabolic dysfunction with neurogenic mechanisms of blood pressure control.

**Key findings:**

- The FGF-21/β-Klotho axis remains unchanged in the CNS of SHRs.
- Hepatic β-Klotho is significantly reduced in hypertensive rats.
- These findings suggest that a peripheral FGF-21 resistance may contribute to the metabolic and inflammatory milieu sustaining hypertension.
- The study highlights the liver as a site linking metabolic dysfunction and neurogenic hypertension.

## Introduction

The central nervous system (CNS) is a major regulator of cardiovascular function, integrating autonomic and endocrine inputs that modulate the blood pressure and heart rate. Within the hypothalamus and brainstem resides neuronal networks that maintain sympathetic and parasympathetic balance (Guyenet, 2006), and their disfunction is critically implicated in the neurogenic hypertension (Paton & Raizada, 2010; Ribeiro et al., 2020; Ribeiro et al., 2015). However, the molecular mediators linking metabolic signals to central autonomic regulation remain incompletely understood.

Fibroblast Growth Factor 21 (FGF-21) has emerged as a key endocrine hormone regulating glucose and lipid metabolism, energy expenditure, and insulin sensitivity (Lewis et al., 2019; Nishimura et al., 2000; Zhang & Li, 2014). Although initially described as a metabolic regulator, recent evidence indicates that FGF-21 also exerts significant effects on cardiovascular function. Nevertheless, these effects remain controversial, as some works report FGF-21 being able to reduce blood pressure (Chen et al., 2020; He et al., 2016), and others that FGF-21 can act on the CNS to induce pressor mechanisms (Santoso et al., 2017; Song et al., 2018). These opposing findings suggest that the cardiovascular effect of FGF-21 depend on the physiological and metabolic context, and may be modulated by tissue-specific sensitivity to the hormone. FGF-21 acts via the fibroblast growth factor receptor 1 (FGFR1) in complex with the obligatory co-receptor β-Klotho (KLB). Impairments in this pathway, particularly reduced β-Klotho expression, have been associated with dysregulated lipid and carbohydrate homeostasis and systemic inflammation, which could be associated to an impairment of the FGF-21 signaling and the establishment of a state of “FGF-21 resistance” (Kobayashi et al., 2016; Somm et al., 2018).

Spontaneously hypertensive rats (SHR) represent a well-established model of neurogenic hypertension, in which sympathetic overactivity is responsible for arterial pressure increases (Judy et al., 1976). Whether the FGF-21 signaling axis is altered during the development of hypertension in this model remains unknown. We hypothesized that hypertension is associated with altered FGF-21/β-Klotho signaling, particularly through a reduction in tissue β-Klotho expression, indicative of peripheral FGF-21 resistance that may contribute to metabolic and autonomic dysregulation. To evaluate this possibility, we investigated the expression of the genes related to the FGF-21 signaling axis (FGF-21, FGFR, and KLB) in the hypothalamus, brainstem and liver of SHR at pre-hypertensive (21 days old - SHR-21d) and hypertensive (90 days old **-** SHR-90d) stages.

## Methodology

### Animals and ethical approval

Male SHR were obtained from the Animal Facility at the Institute of Biomedical Sciences, University of São Paulo, at two different ages: Twenty-one days old (SHR-21d; n=9) and 90 days old (SHR-90d; n=16). Animals were housed at temperatures of 22±2°C with 12/12h light/dark cycle, air humidity of 50–60% with free access to standard rat chow (Nuvilab®, Paulínia, SP, Brazil) and tap water. The animals were used in accordance with the certificates approved by the Ethical Committee for Animal Research of the Institute of Biomedical Sciences, University of São Paulo (CEUA #9048200821), and followed the Ethical Principles in Animal Research issued by the Brazilian College of Animal Experimentation.

### Monitoring of blood pressure and heart rate

Systolic tail pressure (STP **—** mmHg) and heart rate (HR **—** bpm) values were obtained from animals by using the technique of tail-cuff plethysmography (BP-2000 Blood Pressure Analysis System™ — Visitech Systems). Animals were acclimated to the procedure in a silent room for 5 days prior to the day of the experiment during 30 min between 10-11 AM. After acclimation, STP and HR were measured for 25 readings in the same room for 4 days.

### Tissue collection

In the final day of cardiovascular measurements, the animals were anaesthetized with 5% isoflurane (Isoforine; Cristália, SP, Brazil) in medicinal oxygen. After confirming deep anesthesia by assessing the absence of withdrawal reflexes to noxious pinching of the paws and tail, the animals were decapitated for brain collection, and a midline laparotomy was performed to access the liver for tissue sampling. The frontal cortex, brainstem, and hypothalamic block were collected, along with the left lobe of the liver. After collecting, samples were stored in -80°C for posterior gene expression analysis using reverse transcription real-time PCR (RT-qPCR) assay.

### RNA extraction and reverse transcription

Tissue samples were homogenized in TRIzol™ Reagent (Invitrogen — Massachusetts, USA), and the Illustra™ RNAspin Mini Isolation Kit was used to isolate total RNA following manufacturer instructions. Quality control was employed using the NanoDrop® 2000 (Thermo Fisher Scientific — Massachusetts, USA) (total RNA must be ≥ 100 ng/μL) and loading GelRed-stained RNA samples into agarose gels and observing for intact 28S and 18S rRNA bands — samples that failed either test were discarded. Reverse transcription was performed on every viable sample using the SuperScript™ kit (Invitrogen — Massachusetts, USA) and a Veriti™ Thermal Cycler (Applied Biosystems — Massachusetts, USA), following the kit manufacturer’s instructions.

### RT-qPCR procedure

The primers designed for the FGF-21, FGFR1, β-Klotho and β-actin genes are disclosed in Table 1. RT-qPCR reactions were performed using the SsoAdvanced™ Universal SYBR® Green Supermix (Bio-Rad Laboratories — California, USA) and the thermal cycler and software Rotor-Gene Q (Qiagen — Hilden, Germany). Gene expression analysis was performed using the 2^-ΔΔCt^ method, as described by Livak and Schmittgen (Livak & Schmittgen, 2001). The expression levels of the target genes were normalized to β-actin, which exhibited the most stable expression across the analyzed tissues and samples.

**Table 1.**
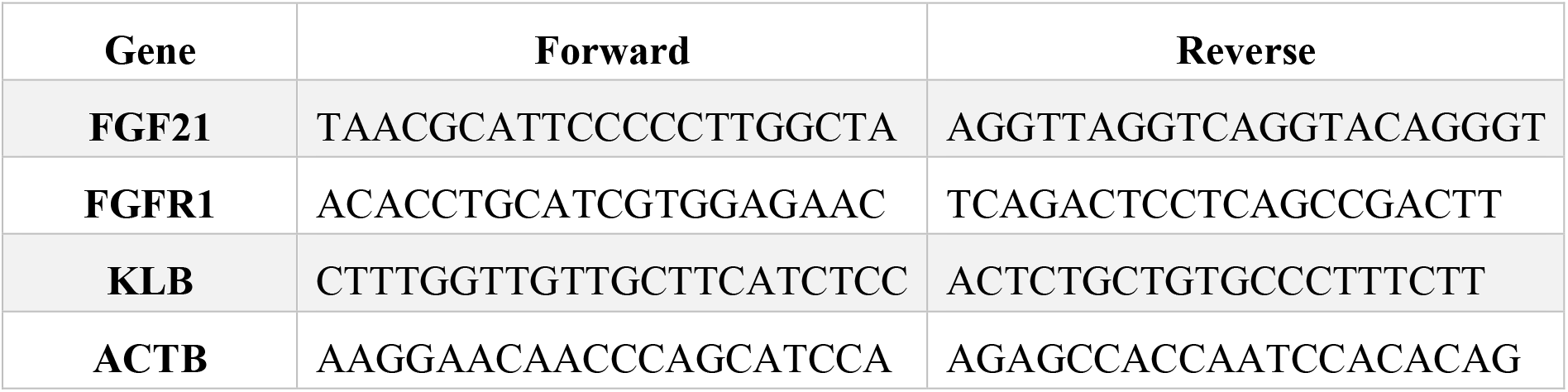
Primers used for quantitative PCR.

### Statistical analysis

Data were analyzed and plotted in the GraphPad Prism 10 ® software (GraphPad Software, Inc. San Diego, CA, USA). Results were expressed as mean ± standard error of mean. Comparisons between groups were conducted using Student’s or Welch’s unpaired *t*-tests after assessing normality and variance of data. Differences were considered statistically significant when p < 0.05.

## Results

### Age-related increase in blood pressure without changes in heart rate in SHR

Figure 1A depicts the absolute values of STP and HR of SHR at different ages showing that SHR-21d presented lower systolic pressure (128±2 mmHg, n=9) when compared to the SHR-90d (177±4 mmHg, n=16, p<0.0001). Figure 1B shows that HR values did not differ between groups (SHR-21d: 412±29 bpm vs. SHR-90d: 402±6 bpm, p=0.1260).

**Figure 1.**
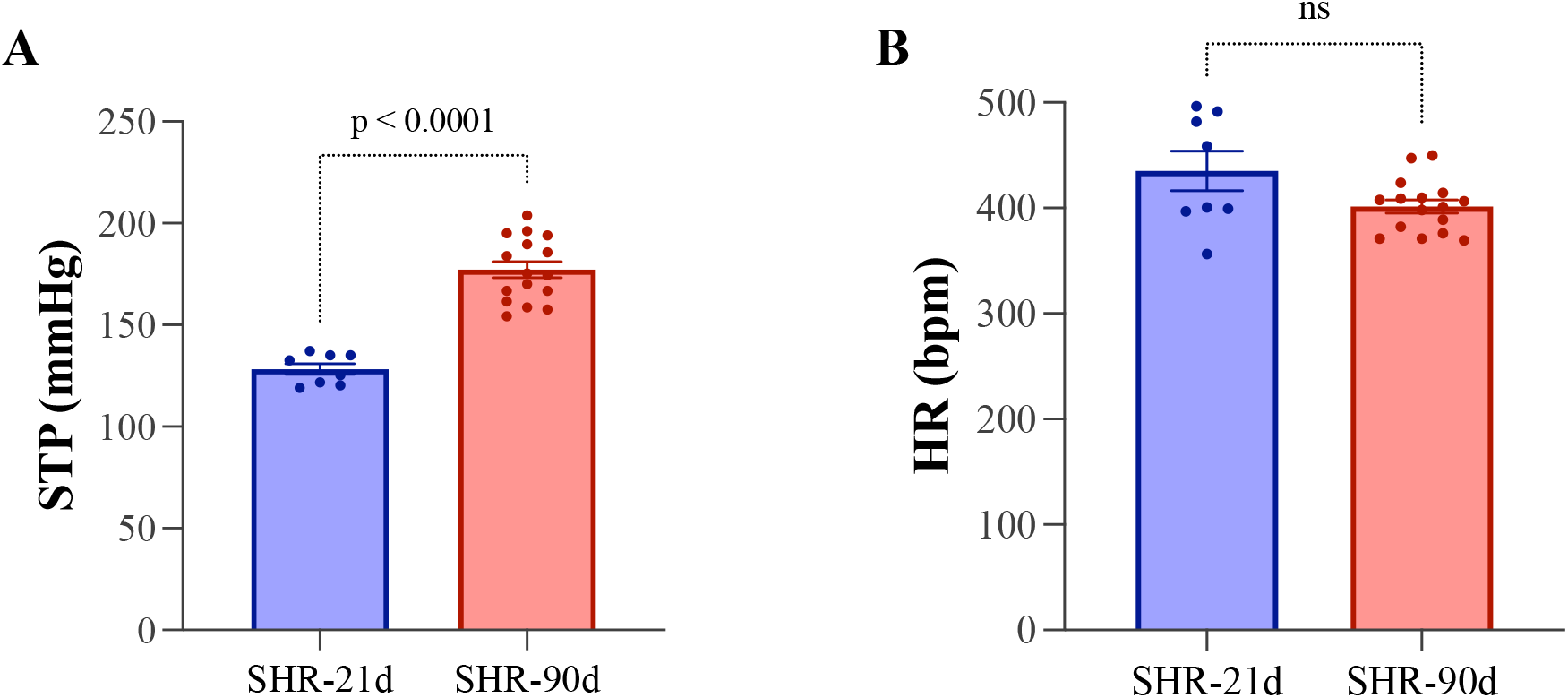
Cardiovascular profile in spontaneously hypertensive rats (SHR). (A) Systolic tail pressure (STP, mmHg) and (B) heart rate (HR, bpm) measured in 21-day-old (SHR-21d) and 90-day-old (SHR-90d) rats. Values are expressed as mean ± SEM. p < 0.0001 vs. SHR-21d.

### Hepatic KLB expression decreases with age in SHR

Relative mRNA expression levels of FGF-21, FGFR1, and KLB were assessed in the liver, brainstem, hypothalamus and frontal cortex of SHR-21d and SHR-90d (Fig 2). Among the tissues analyzed, the liver exhibited the most pronounced differences, with KLB expression significantly lower in SHR-90d compared with SHR-21d (p < 0.05; Fig. 2A), while FGF-21 and FGFR1 transcript levels remained unchanged. In contrast, no significant differences in the expression of any of these genes were detected in the brainstem (Fig. 2B), hypothalamus (Fig. 2C), or frontal cortex (Fig 2D).

**Figure 2.**
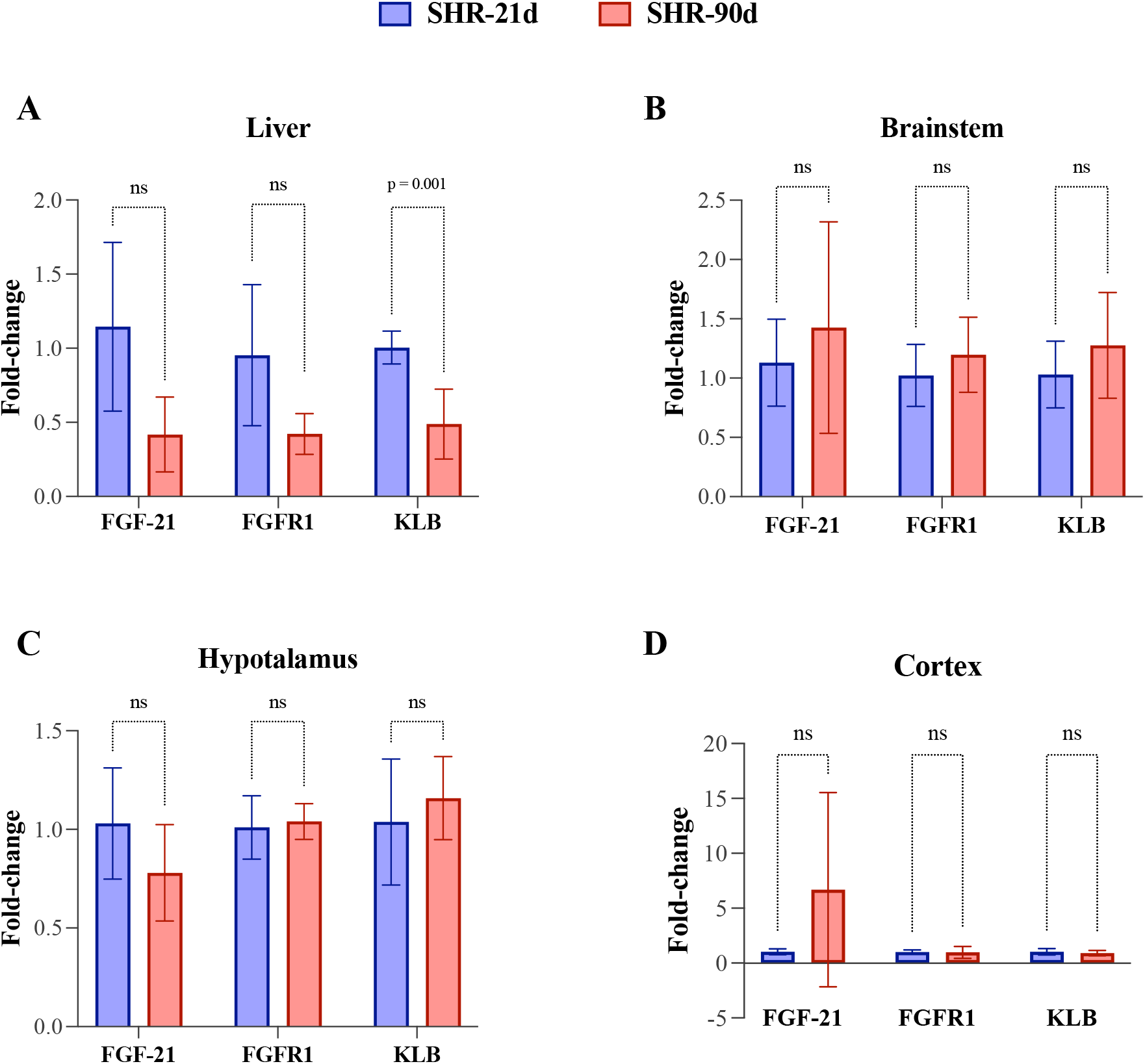
Relative mRNA expression of FGF-21 pathway genes in peripheral and central tissues of SHR. Quantitative PCR analysis of FGF-21, FGFR1, and β-Klotho (KLB) mRNA in the (A) liver, (B) brainstem, (C) hypothalamus, and (D) frontal cortex of SHR at 21 and 90 days of age. Data are normalized to β-actin and expressed as mean ± SEM. p < 0.05 vs. SHR-21d.

### Discussion

This study demonstrates that hypertension in SHR is accompanied by a selective downregulation of hepatic β-Klotho, without significant alterations in the CNS expression of FGF-21, FGFR1, or β-Klotho. These findings suggest a state of peripheral FGF-21 resistance rather than central impairment, providing evidence that hepatic dysregulation of the FGF-21/β-Klotho axis may act as a metabolic signature of the hypertensive phenotype.

FGF-21 is an endocrine fibroblast growth factor predominantly secreted by the liver, acting as a key regulator of lipid and glucose metabolism, insulin sensitivity, and energy expenditure (Kharitonenkov et al., 2005; Markan et al., 2017). Its biological activity depends on binding to fibroblast growth factor receptor 1 (FGFR1) in complex with the co-receptor β-Klotho, which confers tissue specificity (Ogawa et al., 2007). Reduced β-Klotho expression has been described in metabolic disorders such as obesity, type 2 diabetes, and non-alcoholic fatty liver disease, leading to FGF-21 resistance, characterized by high circulating FGF-21 levels and diminished target responsiveness (Fisher & Maratos-Flier, 2016; Tezze et al., 2019).

In the present study, the decrease in hepatic β-Klotho expression in hypertensive SHR (Fig. 2A) likely reflects a similar mechanism of hormone resistance. This hepatic FGF-21 insensitivity could contribute to systemic metabolic dysregulation frequently observed in hypertension, including insulin resistance, altered glucose, lipid and amino-acid metabolism, and low-grade inflammation (da Silva et al., 2020; Li et al., 2023; Makuch-Martins et al., 2024). Importantly, the liver plays a pivotal role in the crosstalk between metabolic and cardiovascular systems, acting through neural, endocrine, and cytokine-mediated pathways (Morton et al., 2014). Thus, hepatic β-Klotho downregulation may attenuate FGF-21 signaling to the CNS, reducing feedback inhibition of sympathetic activity and facilitating the persistence of neurogenic hypertension, a possibility that deserves to be investigated.

Although FGF-21 is primarily known for its metabolic actions, emerging evidence indicates that it also influences cardiovascular function and autonomic regulation. For instance, central or systemic administration of recombinant FGF-21 can lower blood pressure and improve baroreflex sensitivity in animal models of metabolic syndrome and hypertension (Chen et al., 2020; He et al., 2016). Conversely, intracerebroventricular injection of FGF-21 has been shown to activate neurons in the paraventricular nucleus of the hypothalamus, event that has been shown to increase sympathetic outflow and blood pressure (Nomura et al., 2019). These apparently contradictory findings support the notion that the cardiovascular effects of FGF-21 depend on the context and tissue-specific sensitivity of the FGF-21/β-Klotho signaling system. Our data reinforce this concept by showing that the central FGF-21 pathway remains intact, while peripheral hepatic impairment coincides with the development of hypertension.

Taken together, our results identify the hepatic FGF-21/β-Klotho axis as a critical peripheral mechanism potentially contributing to the metabolic–autonomic coupling underlying hypertension. Further studies assessing circulating FGF-21 levels and hepatic receptor sensitivity, as well as interventions restoring β-Klotho expression, will be essential to determine the causal relationship between hepatic FGF-21 resistance and the hypertensive phenotype.

## Acknowledgments

We wish to thank Dr Maristela M. Okamoto for helpful technical support, and the financial support by 1) Sao Paulo State Research Foundation (FAPESP) # 2023/11230-9 (PSS), #2023/08762-9 (VRA); 2) CAPES - Finance Code 001; 3) (GAO) - National Council for Scientific and Technological Development [CNPq – 103575/2023-5]. Antunes VR is CNPq Research Fellow: # 305570/2023-4

